# Stimulus-specific enhancement of responses in mouse primary visual cortex mediated by GABA release from VIP cells

**DOI:** 10.1101/2023.06.19.545641

**Authors:** Megumi Kaneko, Mahmood S. Hoseini, James A Waschek, Michael P. Stryker

## Abstract

When adult mice are repeatedly exposed to a particular visual stimulus for as little as one hour per day for several days while their visual cortex (V1) is in the high-gain state produced by locomotion, that specific stimulus elicits much stronger responses in V1 neurons for the following several weeks, even when measured in anesthetized animals. Such stimulus-specific enhancement (SSE) is not seen if locomotion is prevented. The effect of locomotion on cortical responses is mediated by vasoactive intestinal peptide (VIP) positive interneurons, which can release both the peptide and the inhibitory neurotransmitter GABA. Here we used genetic ablation to determine which of those molecules secreted by VIP-ergic neurons is responsible for SSE. SSE was not impaired by VIP deletion but was prevented by compromising release of GABA from VIP cells. This finding suggests that SSE may result from Hebbian mechanisms that remain present in adult V1.

**SIGNIFICANCE:** Many neurons package and release a peptide along with a conventional neurotransmitter. The conventional view is that such peptides exert late, slow effects on plasticity. We studied a form of cortical plasticity that depends on the activity of neurons that express both vasoactive intestinal peptide (VIP) and the inhibitory neurotransmitter GABA. GABA release accounted for their action on plasticity, with no effect of deleting the peptide on this phenomenon.

## INTRODUCTION

Recordings from alert mice revealed that behavioral state has a dramatic influence on the responses of neurons in the primary visual cortex (Niell and Stryker, 2010). Locomotion, in particular, produced a highly reproducible increase in cortical responses while having little effect on the activity of neurons in the lateral geniculate nucleus (LGN). V1 neurons remained as selective during locomotion as when animals were still, and orientation selectivity was similar to that found in anesthetized animals (Niell and Stryker, 2008).

Vasoactive intestinal peptide (VIP) positive interneurons play a crucial role in the increased responsiveness of V1 neurons during locomotion. When the VIP cells in a small region of V1 were destroyed, V1 responses remained normally selective but were no longer increased by locomotion (Fu et al., 2014, 2015). When VIP cells in V1 were activated optogenetically in alert mice that were still, responses were increased just as if the animals were running (Fu et al., 2014) These findings showed that VIP-cell activity was both necessary and sufficient for the increase in selective V1 responses.

Locomotion also enhanced plasticity in a mouse model of recovery from deprivation amblyopia (Kaneko and Stryker, 2014). When binocular vision was restored in mice that had been deprived of vision in one eye by unilateral lid suture for 4 months beginning in early life, recovery assayed by intrinsic signal imaging was meager, less than halfway to normal levels of response. When these mice were instead given 4 hr/day of visual stimulation during locomotion, the response to the stimuli viewed recovered to normal levels after 5-10 days. Microelectrode recordings confirmed the loss and restoration of visual responses (Kaneko and Stryker, 2014). Blocking synaptic release from the VIP cells by their expression of tetanus toxin light chain prevented recovery of responses, while optogenetic stimulation of VIP cells in stationary mice produced recovery (Fu et al., 2015).

In normal adult mice, viewing a particular stimulus, such as a drifting bar or grating of a particular orientation, for 1 hr/day while running on a polystyrene ball floating on an air stream produced a stimulus-specific enhancement (SSE) of V1 responses to that stimulus. Such SSE lasts for weeks and is evident both with intrinsic signal imaging through an intact skull, which is a completely non-invasive assay, and with single cell measurements (Kaneko et al., 2017). Neurons selective for the stimulus viewed during locomotion increase their responses to it, while the responses of neurons that did not initially respond well to that stimulus are unchanged. Indeed, the resulting enhancement is proportional to the degree to which each neuron responded to the stimulus prior to the exposure during locomotion. Identical exposure to the stimulus without the opportunity for locomotion produced no enhancement (Kaneko et al., 2017).

These findings taken together suggest that some product of VIP-cell activity is responsible for SSE. VIP cells secrete the inhibitory neurotransmitter GABA, which appears principally to inhibit somatostatin-positive (SST) neurons and thereby disinhibit the excitatory neurons of V1. This disinhibition accounts for the increase in responses during locomotion (Fu et al., 2014). VIP cells also secrete the peptide VIP, the effects of which on the targeted cortical cells are not known. The present experiment seeks to determine which of these two VIP-cell secreted molecules is responsible for persistent SSE. We used both intrinsic signal imaging and microelectrode recording to study 3 groups of mice: those in which the peptide VIP was genetically deleted (*Vip*^-/-^); mice in which GABA release from VIP-cells was compromised by knocking out the vesicular GABA transporter (*Vip-Cre;Vgat*^*-/-*^); and control mice, including littermates of the VIP^-/-^ animals. Our conclusion is that SSE depends on GABA release from VIP cells but not on the secretion of the peptide VIP.

## RESULTS

### Characterization of visual responses in V1 of VIP null animals and mice with disrupted Vgat in VIP cells using intrinsic signal imaging

VIP-null (*Vip*^*-/-*^) and mice wildtype at the VIP locus (*Vip*^*+/+*^) were produced by crossing a *Vip*^*+/-*^ female with a *Vip*^*+/-*^ male for the experiments on SSE using intrinsic-signal imaging shown in Figures 1, 3, and 4. To examine the role of vesicular GABA release from VIP cells in SSE, we used the conditional vesicular GABA transporter (*Vgat*) loss-of-function mice, and generated animals for study by crossing *Vgat*^*flox/flox*^ mice with *Vip-Cre::Vgat*^*flox/+*^ mice. The products of this breeding strategy, *Vgat*^*+/-*^ and *Vgat*^*-/-*^ mice were used as control and experimental animals, respectively. Vgat is required for the loading of GABA into synaptic vesicles and therefore essential for the normal synaptic release of GABA. It is well established that synaptic GABA release is absent in cells of the floxed *Vgat* mouse line that express cre recombinase (Wu et al., 2000; Wojcik et al., 2006; Lin et al., 2008; Tong et al., 2008; Saito et al., 2010; Vong et al., 2011; Pei et al., 2015; Hirano et al., 2016).

**Figure 1.**
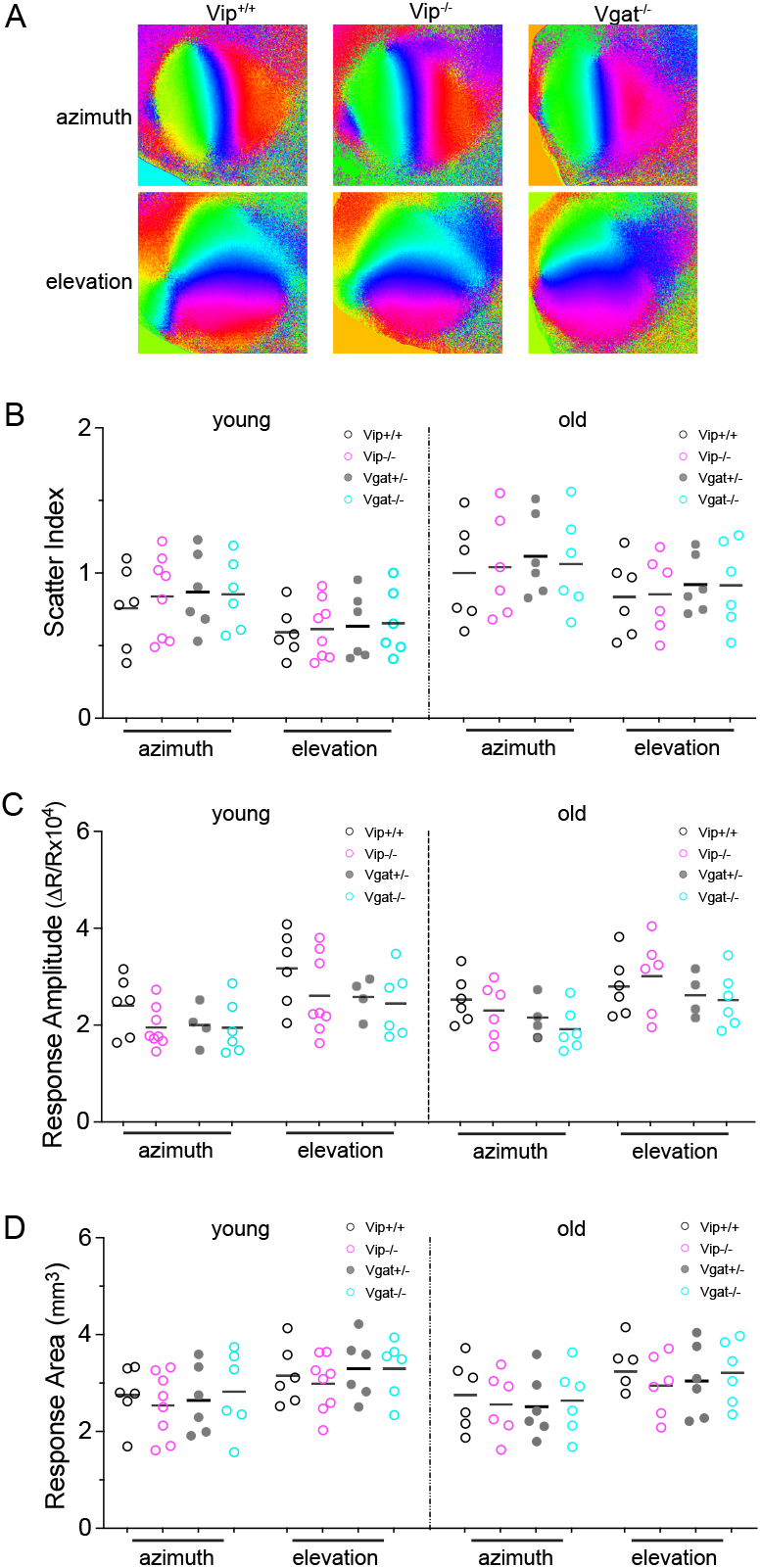
Baseline intrinsic signal responses in Vip^-/-^ and Vgat^-/-^ animals are indistinguishable from those in control animals. A. Examples of retinotopic maps recorded with intrinsic signal imaging, in Vip^+/+^, Vip^-/-^, and Vgat^-/-^ mice at young adult stage (P120 ∼ 140). Azimuth and elevation maps are generated from responses to drifting vertical bars and horizontal bars, respectively. B -D. Map statistics from Vip^-/-^ and Vgat^-/-^ experimental mice and Vip^+/+^ and Vgat^+/-^ controls. B. Map scatter index. C. Response amplitude. D. Response area. Left panels in B - D: Young adults (P120 ∼ 140), and right panels in B - D: Old mice (P210 ∼ 240). Data are statistically indistinguishable among Vip^+/+^, Vip^-/-^, Vgat^+/-^, and Vgat^-/-^ animals, both in young and old adults.

We first examined visual cortical responsiveness at a global level using intrinsic signal imaging (Kalatsky and Stryker, 2003; Cang et al., 2005) under isoflurane anesthesia in young adult mice (P120 – 140) and in old mice (P210 –240) (Figure 1). Examples of cortical retinotopic maps in response to vertical bars drifting right – left (azimuth maps) and to horizontal bars drifting up – down (elevation maps) are shown in Figure 1A. The gross polarity of the retinotopic map in V1 was largely normal in both Vip^-/-^ and Vgat^-/-^ mice. To compare the maps quantitatively, we computed the “map scatter” by calculating the differences between the phase values of the individual pixels within the visual area to those of their near neighbors, as previously described (Cang et al., 2005). For maps with high quality and strong visual response, these phase differences should be quite small due to the smooth progression of V1 topography. For the azimuth maps, the map scatter in Vip^+/+^ mice was 0.75 ± 0.28º (young, n = 6) and 1.0 ± 0.35º (old, n = 6), in Vip^-/-^ mice 0.84 ± 0.29º (young, n = 8) and 1.04 ± 0.35º (old, n = 6), in Vgat^+/-^ mice 0.87 ± 0.27º (young, n = 6) and 1.12 ± 0.28º (old, n = 6), and in Vgat^-/-^ mice 0.85 ± 0.25º (young, n = 6) and 1.06 ± 0.33º (old, n = 6); all statistically insignificant from each other (figure 1B). Scatter index in elevation maps are similarly comparable among Vip^+/+^,Vip^-/-^, Vgat^+/-^, and Vgat^-/-^ animals. Vip^+/+^: 0.59 ± 0.17º (young), 0.83 ± 0.27º (old); Vip^-/-^0.62 ± 0.20º (young), 0.85 ± 0.27º (old); Vgat^+/-^: 0.63 ± 0.23º (young), 0.92 ± 0.20º (old); Vgat^-/-^ : 0.66 ± 0.23º (young), 0.92 ± 0.30º (old) (Figure 1B).

Cortical responses to visual stimulation in Vip^-/-^ mice and Vgat^-/-^ mice were not statistically different in amplitude from those in Vip^+/+^ and Vgat^+/-^ animals, respectively, at a global level as measured by intrinsic signal optical imaging (Figure 1C). The average amplitude of azimuth maps in Vip^+/+^ mice: 2.4 ± 0.61 deltaR/R (young), 2.53 ± 0.49 (old); in Vip^-/-^ mice: 1.95 ± 0.42 (young), 2.3 ± 0.56 (old); in Vgat^+/-^ mice: 2.00 ± 0.43 (young), 2.16 ± 0.42 (old); in Vgat^-/-^ mice: 1.95 ± 0.42 (young), 1.92 ± 0.45 (old). The average amplitude of horizontal maps in Vip^+/+^ mice: 3.17 ± 0.78 (young), 2.85 ± 0.61 (old); in Vip^-/-^ mice: 2.61 ± 0.82 (young), 3.01 ± 0.78 (old); in Vip^+/-^ mice: 2.58 ± 0.41 (young), 2.62 ± 0.46 (old); in Vgat^-/-^ mice: 2.45 ± 0.69 (young), 2.52 ± 0.58 (old).

Similarly, the response areas in both Vip^-/-^ mice and Vgat^-/-^ mice were indistinguishable from those in respective control animals, for both azimuth and elevation maps in both young age and old adult mice (Fig 1D).

These observations indicate that, with normal visual experience, visual cortical function in Vip^-/-^ mice and Vgat^-/-^ mice develop without detectable impairment in the measures we evaluated.

### Response properties and selectivity of single neurons

For the microelectrode recording experiments shown in Figure 2, C57BL/6 mice were purchased from Jax and bred for use as wildtype control animals. Microelectrode recordings made in alert *Vip*^*-/-*^ and *Vip-Cre::Vgat*^*-/-*^ mice were compared to those made in WT mice in terms of firing rates to gratings, indices of ocular dominance and orientation selectivity, and variability in response. Head-fixed mice viewing a video monitor were free to stand or run on a polystyrene ball floating on air while 2-shank silicon probes with 128 recording sites were lowered into the binocular zone of V1. After spike sorting, the waveforms of isolated single units were classified as broad-spiking (putatively excitatory) or narrow-spiking (putatively inhibitory). Mice in these experiments had no prior practice on the polystyrene ball and were therefore mostly still, rather than walking or running.

**Figure 2.**
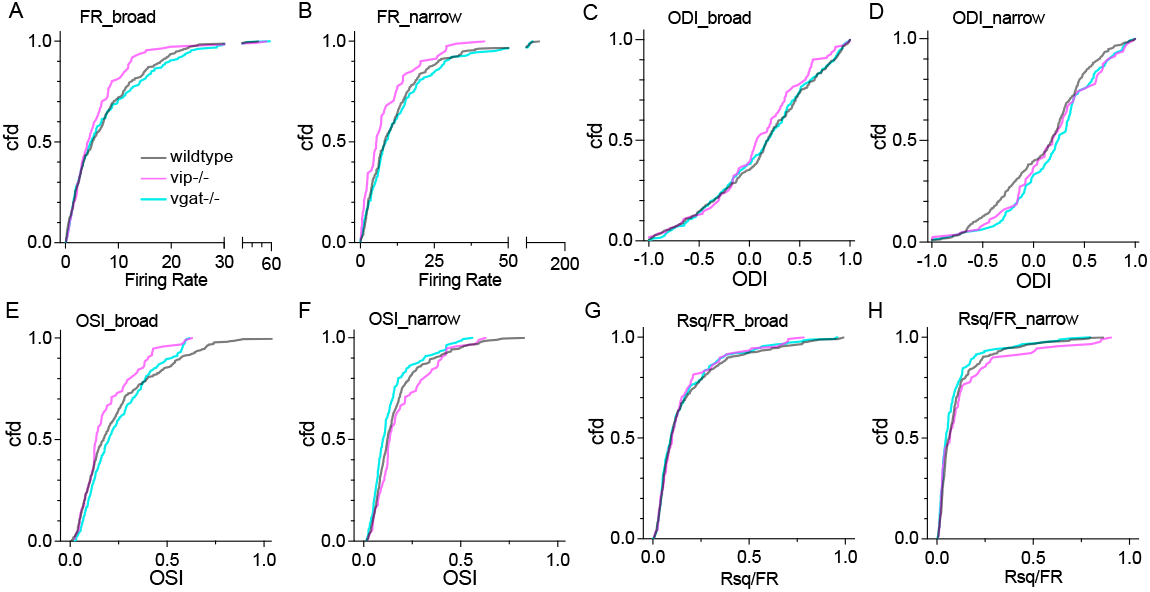
Response properties in individual neurons in V1 examined by single unit electrophysiological recording A, B. Cumulative frequency distribution of firing rate of broad-spiking cells (A) and narrow-spiking cells (B) in response to contralateral-eye stimulation during still condition. C, D. Cumulative frequency distribution of ocular dominance index in broad-spiking cells (C) and narrow-spiking cells (D). E, F. Cumulative frequency distribution of orientation selectivity index in broad-spiking cells (E) and narrow-spiking cells (F). G, H. Cumulative frequency distribution of a measure of response variability. Data from WT, Vip^-/-^, and Vgat^-/-^ animals are presented in grey, magenta, and cyan lines, respectively. Sample size in A - H: WT, broad-spiking: 374, narrow-spiking: 295; Vip^-/-^, broad-spiking: 115, narrow-spiking: 81; Vgat^-/-^ broad-spiking: 262, narrow-spiking: 167.

**Figure 3.**
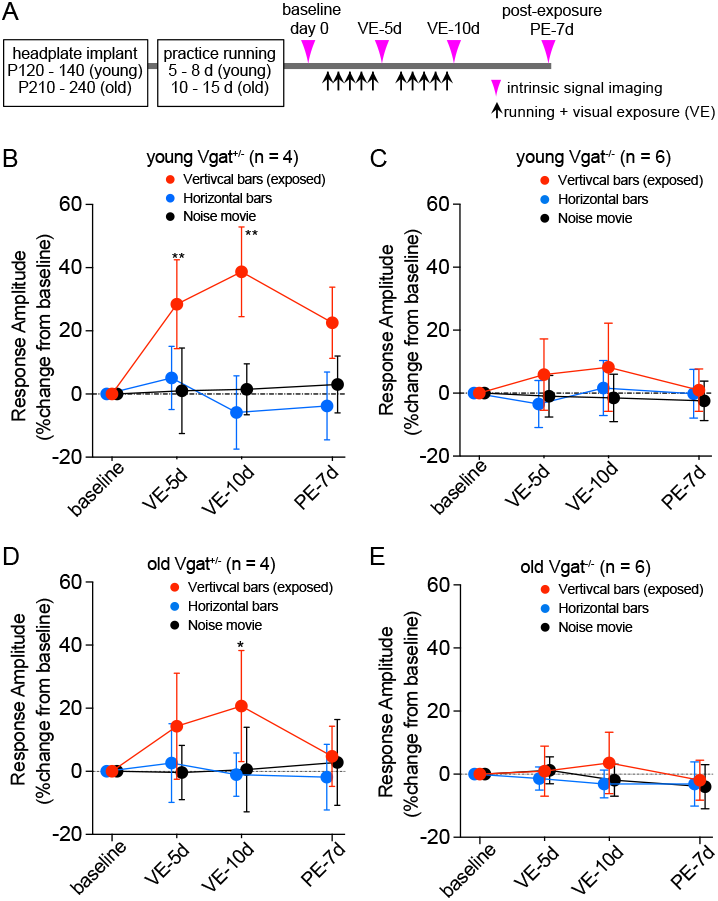
Stimulus-specific response enhancement is impaired in mice with Vgat loss-of-function. A. Experimental schedule. B - C. Change in response magnitude in young adult mice to exposed stimuli (vertical bars, red circle) and unexposed stimuli (horizontal bars, blue circle, contrast-modulated noise movie, black circle) in control (Vgat^+/-^, B) and experimental animals (Vgat^-/-^, C). D - E. Change in response magnitude in old mice to exposed stimuli (vertical bars, red circle) and unexposed stimuli (horizontal bars, blue circle; contrast-modulated noise movie: black circle) in Vgat^+/-^ control animals (D) and in Vgat^-/-^ experimental animals (E). Response amplitude is expressed as percent change from baseline (100x[post - baseline]/baseline). Error bars represent mean ± s.d. **P<0.01, *P<0.05 vs. baseline response to each stimulus; two-way ANOVA followed by multiple comparisons with Bonferroni corrections.

**Figure 4.**
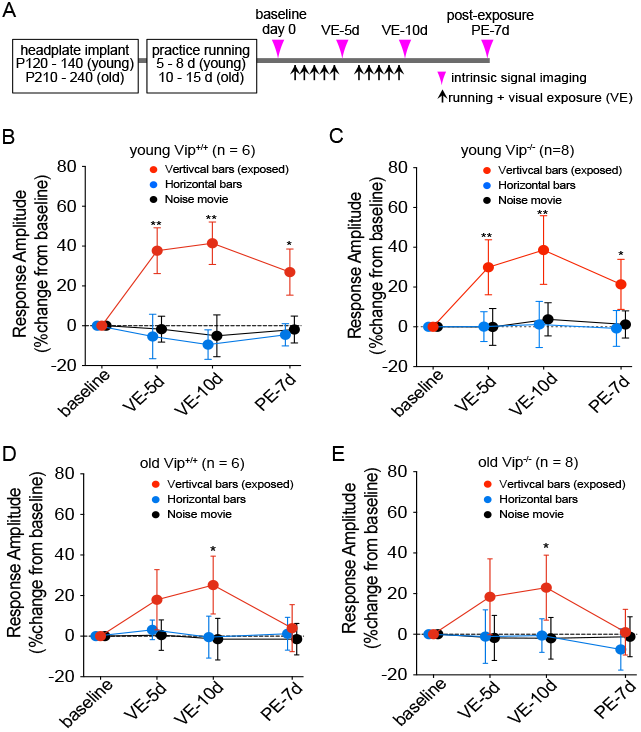
Stimulus-specific enhancement in VIP null mice is indistinguishable from that in Vip^+/+^ control mice. A. Experimental schedule. B - C. Change in response magnitude during young adult age to exposed stimuli (vertical bars: red circle) and unexposed stimuli (horizontal bars: blue circle, contrast-modulated noise movie: black circle) in Vip^+/+^ control animals (B) and in VIP-null mice (C). D - E. Change in response magnitude in old mice to exposed stimuli (vertical bars: red circle) and unexposed stimuli (horizontal bars: blue circle, contrast-modulated noise movie: black circle) in Vip^+/+^ control animals (D) and in VIP-null mice (E). Response amplitude is expressed as percent change from baseline (100x[post - baseline]/ baseline). Error bars represent mean ± s.d. **P<0.01, *P<0.05 vs. baseline response to each stimulus.

Figure 2 shows cumulative frequency distributions for each of the measures above. Because firing rates are strongly influenced by locomotion, we compare them in Figure 2A,B only during comparable still periods. Firing rates elicited by stimulation through the contralateral eye of both broad- and narrow-spiking neurons were essentially identical in Vgat^-/-^ and WT mice but both appeared slightly lower in Vip^-/-^ mice, although not significantly so (Figure 2A, B). The ocular dominance distributions of both broad- and narrow-spiking neurons were nearly identical in the 3 groups of mice (Figure two experimental groups of mice were similar to and 2C, D). The distributions of orientation selectivity as measured through the contralateral eye were not significantly different in the 3 groups. As expected, the broad-spiking neurons (Figure 2E) were much more selective than the narrow-spiking neurons (Figure 2F) in all 3 groups. We constructed a measure of response variability for each neuron analogous to coefficient of variation by dividing the “goodness of fit” of the orientation tuning curve by the neuron’s firing rate at its optimal orientation. These distributions of response variability, which exclude periods of locomotion, were nearly identical in the 3 groups of mice for both broad- and narrow-spiking neurons (Figure 2G, H).

Before the critical period of susceptibility to the effects of monocular visual deprivation, at around 4 weeks of age, the preferred stimulus orientation of individual V1 neurons driven through one eye is only randomly related to the preferred orientation measured through the fellow eye (Wang et al., 2010). After visual experience during the critical period, orientation selectivity measured through the two eyes becomes similar. One measure of the normal development of visual cortical organization, therefore, is the similarity of preferred orientation between orientation-selective neurons as assessed through the two eyes. The median differences in orientation selectivity for the statistically indistinguishable from those of WT mice (medians and interquartile range: Vip-Vgat^-/-^ 29°±23°, Vip^-/-^ 26°±22°, WT 24°±16°).

### Vesicular GABA release from VIP cells is required for stimulus-specific response enhancement

After confirming that cortical responsiveness to visual stimuli in Vgat^-/-^ and Vip^-/-^ mice is largely normal at a global level, we next tested whether disruption of GABAergic transmission from VIP cells affects SSE. Figure 3A shows the experimental schedule. Each animal was acclimated to the experimental apparatus by handling and being placed on the polystyrene ball floating on air in a dimly lighted room with no moving visual stimuli. Young adult mice were acclimated for approximately 15 min a day for 5 – 8 days, and old mice for 10 – 15 days. Baseline intrinsic signal responses were then measured to three different, high-contrast visual stimuli: a drifting horizontal bar, a drifting vertical bar, and contrast-modulated noise movie. Starting 2 – 3 days later, animals were allowed to run on polystyrene balls while viewing the drifting vertical bar for 60 min per day for 10 days, as described previously (Kaneko et al., 2017). Our earlier experiments had shown that the effect of 1 hr/day of exposure was saturating; doubling the exposure time per day did not increase or accelerate SSE (Kaneko et al., 2017). To track the change in visual cortical responses to these visual patterns, we repeated intrinsic signal recordings on days 5 and 10; and additionally on day 17, one week after stopping the daily running + visual exposure (VE) sessions (Figure 3A).

Young adult Vgat^+/-^ control mice showed a significant increase in response only to the vertical bar that the animal viewed during daily running (28.4 ± 14.1%, red circles in Figure 3B), while responses to other stimuli were unchanged (horizontal bar: -5.8 ± 11.6%, blue circles; noise movie: 1.0 ± 13.6%, black circles). This enhancement had nearly peaked by day 5 with only a slight increase on day 10 (38.7 ± 14.2%) and it persisted for at least 7 days after terminating sessions of running + VE (22.6 ± 11.3%) (Figure 3B).

Significant response enhancement to the exposed stimulus in old Vgat^+/-^ control mice was apparent only after 10 days of running + VE, and was smaller than that in young mice (14.3 ± 16.8% at 5d, 20.1 ± 17.6% at 10d) (Figure 3D). This small enhancement in old heterozygous control mice did not persist 7 days post-exposure (4.8 ± 9.6%; Figure 3D). The current observations in the young heterozygous control mice are consistent with our previous report (Kaneko et al., 2017).

In contrast to the heterozygous control animals, Vgat^-/-^ mice failed to show enhancement of responses to the stimulus that the animal viewed during daily running. This impairment was observed both in young adult mice (5d: 8.4 ± 16.7%, 10d: 8.1± 8.3%; both P>0.99 vs. baseline; Figure 3C) and old mice (5d: 1.0 ± 3.6%, 10d: 7.9 ± 9.7%; both P>0.99 vs. baseline, Figure 3E).

### VIP is not required for stimulus-specific response enhancement

Mice deficient in VIP (Vip^-/-^) have previously been reported to exhibit various neurological, developmental, and/or behavioral deficits (Colwell et al., 2003; Harmar, 2003; Itri et al., 2004; Aton et al., 2005; Loh et al., 2014). We first examined V1 visual responses in Vip^-/-^ mice at a global level, using intrinsic-signal imaging to measure retinotopic organization and response magnitude, as shown in Figure 1. After confirming largely normal visual cortical responsiveness of Vip^-/-^ mice, we next tested whether the absence of VIP affects SSE. Figure 4A shows the experimental schedule. Consistent with our previous report (Kaneko et al., 2017) and with our findings above in Vgat^+/+^ mice, young adult Vip^+/+^ control mice showed a significant increase in cortical responses to the exposed stimulus (vertical bar) after daily visual exposure while running (5d: 37.8 ± 11.6%; 10d: 41.5 ± 10.7%), which persisted at 7d post-exposure (27 ± 11.6%), while no significant change to unexposed stimuli was detected (horizontal bar 5d: -5.4 ± 11.2%, 10d: -9.5 ± 7.4%; noise movie 5d: -1.7 ± 6.5%, 10d: -5.1 ± 10.5%) (Figure 4B). Similarly, a small but statistically significant increase in response to the exposed stimulus was observed in old Vip^+/+^ animals after 10 days of daily running + VE (5d: 17.9 ± 14.9%, 10d: 25.3 ± 14.3%), which did not persist 7d post-exposure (4.0 ± 11.5%, Figure 4D).

In Vip^-/-^ mice, the increase in response magnitude to the exposed stimulus during VE was statistically indistinguishable from that in WT control mice, both in young adults (5d: 29.9 ± 13.8%, 10d: 38.6 ± 17.2%) and in old animals (5d: 16.4 ± 7.6%; 10d: 22.9 ± 6.5%). Persistence of enhancement was also similar to Vip^+/+^ control animals both in young adults and old mice (7d post-exposure: 21.4 ± 12.6% and 1.0 ± 4.6%, respectively) (Figure 4C, E).

## DISCUSSION

### Development of visual cortex with mutant VIP cells

The present experiments reveal that visual cortical organization developed normally when VIP cells were compromised either by the deletion of the VIP peptide or by the impairment of GABA release. The maps of the visual cortex obtained by intrinsic signal imaging had normal orientation and signal-to-noise in the two genetically compromised mouse strains, and the magnitudes of visual responses measured in intrinsic signal imaging were also like those in WT mice. In addition, microelectrode recordings of the responses of single neurons revealed highly similar distributions of all measures of visual response that were obtained. We conclude from this that VIP cells need not release either GABA or VIP peptide for the normal development of these functions of the visual cortex.

### VIP cells in adult plasticity

Adult plasticity is a different matter. Because of our earlier findings on the requirement of locomotion for SSE, together with the requirement for synaptic release from VIP cells for enhancing responses during locomotion, synaptic release from VIP cells was hypothesized to be necessary for SSE. It should be noted that SSE is distinct from the response enhancement during locomotion. Locomotion enhances the responses of all neurons when stimulated appropriately and increases the information about the visual scene carried by activity in V1, but these effects cease when the mouse stops running (Dadarlat and Stryker, 2017). SSE is a persistent enhancement of response to the particular stimulus that was presented during locomotion, and the enhancement is present no matter how the responses are measured or in what state, whether non-invasively in anesthetized animals using intrinsic signal imaging through an intact skull, using 2-photon calcium imaging to track responses of single cells longitudinally, or in alert animals using extracellular microelectrode recording (Kaneko et al., 2017).

While synaptic release from VIP cells was hypothesized to be essential for SSE, previous findings left open the question: release of what? VIP cells release both GABA and VIP peptide, which could act as a neuromodulator. Our previous manipulations of VIP cells--ablation and tetanus toxin expression--similarly affect both products. We therefore sought to eliminate them one at a time to assay their separate effects on SSE. Lacking a conditional knockout of VIP peptide, we were justified in interpreting results in V1 of the global knockout of VIP peptide by the fact that the peptide is expressed nowhere in the brain other than in VIP cells. A conditional knockout of the vesicular GABA transporter allowed us to delete it specifically in VIP cells.

### Relation to other reports of similar plasticity

Previous studies using evoked potential recordings through electrodes implanted in the visual cortex reported a rapid and persistent increase in layer 4 responses to repeated stimuli that was similar to SSE in magnitude and dependence on NMDA-receptor signaling (Frenkel et al., 2006; Cooke and Bear, 2010). However, the increase seen with evoked potentials differed from SSE in that it was found even in restrained animals and is therefore unlikely to depend on VIP cell activity. Possible reasons for these differences are discussed in (Kaneko et al, 2017).

Another form of cortical plasticity depends not on VIP cells but on interneurons derived from the medial ganglionic eminence, principally parvalbumin- and somatostatin-positive interneurons (Southwell et al., 2014). Transplantation of embryonic precursors of either of these types of interneurons into postnatal visual cortex induces a second critical period of ocular dominance plasticity after the end of the normal critical period (Southwell et al., 2010). Interestingly, deletion of Vgat from the transplanted interneurons completely blocks this form of plasticity as well (Hoseini et al., 2019; Priya et al., 2019).

### Conclusion

The findings of the present experiment were unequivocal. Compromising GABA release from VIP cells completely blocked SSE, whereas blocking the release of VIP peptide had no effect on this form of plasticity. The effects were consistent in both young adult and old adult mice. These findings suggest that the effect of VIP cell activity on the firing of the other neurons of the visual cortical circuit is responsible for SSE. Since VIP cell activity inhibits SST cells and thereby disinhibits the excitatory neurons, it allows them to respond more vigorously to the visual stimuli for which they are selective. SSE might then result from Hebbian plasticity mechanisms in V1 that could strengthen the inputs to the excitatory neurons in proportion to the correlation between pre- and postsynaptic activity. Stimulating with a bar of a particular orientation while the activities of the V1 neurons that respond to it are elevated should engage whatever Hebbian plasticity mechanisms are present to make those specific inputs more effective and those responses persistently stronger.

## MATERIALS AND METHODS

### Animals and surgery

All procedures were approved by the Institutional Animal Care and Use Committee of University of California San Francisco. Vgat^fl/fl^ (catalog # 012897), VIP-IRES-Cre knock-in (*Vip*^*tm1(cre)Zjh*^/J; catalog #010908) (Taniguchi et al., 2011), and C57BL/6J (JAX 00664) wild type mice breeders were purchased from Jackson Laboratory (Bar Harbor, ME) and bred as needed, and animals of either sex were used. VIP-null mice were generated as previously described (Colwell et al., 2003) and maintained by Dr. Waschek’s laboratory at University of California Los Angeles. VIP-null (*Vip*^*-/-*^) and mice wildtype at the *VIP* locus (*Vip*^*+/+*^) were produced by crossing a *Vip*^*+/-*^ female with a *Vip*^*+/-*^ male for the experiments on SSE using intrinsic signal imaging shown in Figures 1, 3, and 4. For the microelectrode recording experiments shown in Figure 2, C57BL/6 mice were purchased from Jax and bred for use as wildtype (*Vip*^*+/+*^) control animals.

A custom stainless steel plate for head fixation was attached to the skull with dental acrylic under isoflurane anesthesia. The exposed surface of the skull was covered with a thin coat of nitrocellulose (New-Skin, Medtech Products Inc., NY) to prevent desiccation, reactive cell growth, and destruction of the bone structure. Animals were given a subcutaneous injection of a NSAID carprofen (5 mg/ kg) just before and one day after the surgery. All mice were housed in groups of 4 – 5 and kept under standard conditions (12-h light/12-h dark cycle, with free access to food and water before the surgery, between recordings, and following daily running on the treadmill.

Because stimulus-specific enhancement declines with advanced age in mice, we studied separate groups of young adult (postnatal day (P)120 - 140) and old (P210 - 240) mice.

### Daily exposure to visual stimuli during locomotion

Each animal was acclimated to the running setting by the experimenter’s handling and being placed on the polystyrene ball floating on air for approximately 15 min a day for 5 – 8 days in young animals or 10 – 15 days in old animals. Two to 3 days after measurement of baseline intrinsic-signal responses, animals were allowed to run on a 20-cm polystyrene ball for 60 min per day for 10 days, while viewing a 2º × 60º vertical bar drifting horizontally in both directions, presented on a 30 × 40 cm LCD monitor placed 25 cm from animal’s right eye in a dark room, as described previously (Kaneko et al., 2017).

### Intrinsic signal optical imaging

Repeated optical imaging of intrinsic signals was performed as described (Kaneko et al., 2008). Five to 7 days after the headplate implantation, the first imaging of intrinsic signals was performed to measure baseline responses. The mouse was anesthetized with isoflurane (3% for induction and 0.7% during recording), supplemented with intramuscular injection of chlorprothixene chloride (2 μg/g body weight), and images were recorded transcranially through the window of the implanted headplate. Intrinsic signal images were obtained with a Dalsa 1M30 CCD camera (Dalsa, Waterloo, Canada) with a 135 mm × 50 mm tandem lens (Nikon Inc., Melville, NY) and red interference filter (610 ± 10 nm). Frames were acquired at a rate of 30 fps, temporally binned by 4 frames, and stored as 512 × 512 pixel images after binning the 1024 × 1024 camera pixels by 2 × 2 pixels spatially. Responses in each mouse were measured with three kinds of visual stimuli (1) horizontal bars drifting upward or downward, (2) vertical bars drifting leftward or rightward, and (3) the contrast-modulated noise movie. They were generated in Matlab using Psychophysics Toolbox extensions (Brainard, 1997; Pelli, 1997), and displayed on an LCD monitor (30 × 40 cm, 600 × 800 pixels, 60-Hz refresh rate) placed 25 cm from the mouse, spanning ∼60º (height) × ∼77º (width) of visual space. The drifting bar was the full length of the monitor and 2º wide, and it moved continuously and periodically (Kalatsky and Stryker 2003). The contrast-modulated Gaussian noise movie consisted of the Fourier-inversion of a randomly generated spatiotemporal spectrum with low-pass spatial and temporal cutoffs applied at 0.05 cpd and 4 Hz, respectively (Niell and Stryker, 2008). To provide contrast modulation, the movie was multiplied by a sinusoid with a 10-s period. Movies were generated at 60 × 60 pixels and then smoothly interpolated by the video card to 480 × 480 to appear ∼60º (height) × ∼60º (width) on the monitor and played at 30 frames per second. Each recording took 240 s and was repeated for at least 6 measurements per animal. During the daily session of running + visual exposure (VE), animals were exposed to only drifting vertical bars as described above.

### Analysis of intrinsic signal images

The ROI within V1 was selected on the response magnitude map evoked by visual stimulation. First, the map was smoothed to reduce pixel shot noise by low-pass filtering using a uniform kernel of 5 × 5 pixels. The background area was selected from the area covering ∼150 × 150 pixels outside of V1. The ROI was selected by thresholding at 300% above the average background amplitude and the response amplitude was then calculated as the average amplitude of pixels within the ROI.

### Microelectrode recording

Extracellular recording was carried out in awake, head-fixed mice as in (Hoseini et al 2019) that were free to run on the floating polystyrene ball. On the day of recording, the animal was anesthetized and a craniotomy of about 1-2 mm in diameter was made above the binocular zone of V1 (identified by intrinsic signal imaging). This small opening was large enough to allow for insertion of a 1.1-mm-long double-shank 128-channel electrode fabricated by the Masmanidis laboratory (Du et al., 2011) and assembled by the Litke laboratory (University of California—Santa Cruz). The electrode was placed at an angle of 20-40° to the cortical surface and inserted to a depth of 500-1000 μm. Recordings were started an hour after implantation. For each animal, the electrode was inserted not more than twice.

Visual stimuli were displayed on an LCD monitor (Dell, 30×40 cm, 60 Hz refresh rate, 32 cd/m^2^ mean luminance) placed 25 cm from the mouse (−20° to +40° elevation). Drifting sinusoidal gratings at 12 evenly spaced directions (30 steps, 1.5 s duration, 0.04 cycles per degree, and 1 Hz temporal frequency) were generated and presented in random sequence using the MATLAB Psychophysics Toolbox followed by 1.5-second blank period of uniform 50% gray. Movement signals of locomotion from optical mice tracking the polystyrene ball were acquired in an event-driven mode at up to 300 Hz, and integrated at 100 ms intervals and then converted to the net physical displacement of the top surface of the ball. A mouse was said to be running on a single trial if its average speed for the first 500 ms of the trial fell above a threshold, found individually for each mouse (1–3 cm/s), depending on the noise levels of the mouse tracker. Data acquisition was performed using an Intan Technologies RHD2000-Series Amplifier Evaluation System, sampled at 20 kHz; recording was triggered by a TTL pulse at the moment visual stimulation began.

Single units were identified using MountainSort (Chung et al., 2017), which allows for automated spike sorting of the data acquired using 128-site electrodes. Following manual curation, typical yields ranged between 50-130 isolated single units. Average waveforms of isolated single units were used to calculate three parameters by which cells were classified into narrow- or broad-spiking.

Orientation tuning curves from which response parameters were calculated were fitted as a mixture of two gaussians of individually variable height and width separated by 180 deg for visual stimulus space.

### Statistical analyses

Data were presented as mean ± SEM or mean ± s.d., unless otherwise indicated. Statistical methods employed are stated in the Result section and/or figure legends. Statistical analyses were performed using Prism (GraphPad Software, CA) or Matlab (MathWorks, MA).

## AUTHOR CONTRIBUTIONS

MK and MPS designed the experiment and wrote the manuscript. MK carried out intrinsic signal imaging experiments and analysis. MSH carried out and analyzed electrophysiology experiments. JAW bred and supplied VIP knockout mice. MK, MPS, MSH and JAW revised the manuscript.

## ACKNOWLEDGEMENTS

Supported by NIH Grant R01EY002874. MPS is recipient of the RPB Disney Award for Amblyopia Research. We are grateful to members of the Stryker lab for critical readings.

